# Poly (I:C)-induced maternal immune activation modifies ventral hippocampal regulation of stress reactivity: prevention by environmental enrichment

**DOI:** 10.1101/2021.01.21.427695

**Authors:** Xin Zhao, Ruqayah Mohammed, Hieu Tran, Mary Erickson, Amanda C. Kentner

**Affiliations:** School of Arts & Sciences, Health Psychology Program, Massachusetts College of Pharmacy and Health Sciences, Boston Massachusetts, United States 02115

**Keywords:** Environmental programming, Maternal immune activation, Prenatal immune activation, Gestational, Poly (I:C), Poly I:C, Toll-like 3 receptors, Inflammation, Fetal programming, Maternal behavior, Maternal care, Corticosterone, Glucocorticoid receptor, Corticotropin releasing hormone receptor, Corticotropin releasing factor, Synaptic plasticity, Sex difference, Social interaction, Social approach, Social vigilance, Experience, Inflammation, Schizophrenia, Autism, Social, Hippocampus, Prefrontal cortex, Suprammamillary nucleus, Hypothalamus, Oxytocin, Vasopressin, cFos, CamkIIa, Protein Kinase

## Abstract

Environmental enrichment (EE) has been successfully implemented in human rehabilitation settings. However, the mechanisms underlying its success are not understood. Incorporating components of EE protocols into our animal models allows for the exploration of these mechanisms and their role in mitigation. Using a mouse model of maternal immune activation (MIA), the present study explored disruptions in social behavior and associated hypothalamic pituitary adrenal (HPA) axis functioning, and whether a supportive environment could prevent these effects. We show that prenatal immune activation of toll-like receptor 3, by the viral mimetic polyinosinic-polycytidylic acid (poly(I:C)), led to disrupted maternal care in that dams built poorer quality nests, an effect corrected by EE housing. Standard housed male and female MIA mice engaged in higher rates of repetitive rearing and had lower levels of social interaction, alongside sex-specific expression of several ventral hippocampal neural stress markers. Moreover, MIA males had delayed recovery of plasma corticosterone in response to a novel social encounter. Enrichment housing, likely mediated by improved maternal care, protected against these MIA-induced effects. We also evaluated *c-Fos* immunoreactivity associated with the novel social experience and found MIA to decrease neural activation in the dentate gyrus. Activation in the hypothalamus was blunted in EE housed animals, suggesting that the putative circuits modulating social behaviors may be different between standard and complex housing environments. These data demonstrate that augmentation of the environment supports parental care and offspring safety/security, which can offset effects of early health adversity by buffering HPA axis dysregulation. Our findings provide further evidence for the viability of EE interventions in maternal and pediatric settings.

**Highlights:**

‐ Environmental enrichment (EE) protocols are used clinically to promote rehabilitation
‐ Use of EE in animal models may identify mechanisms underlying clinical successes
‐ Maternal immune activation (MIA) decreased social engagement; this effect was blocked by EE
‐ MIA reduced *c-Fos* activation in the dentate gyrus, while EE reduced activation in the hypothalamus, in response to social stimuli
‐ EE inhibited MIA-induced HPA dysregulation in ventral hippocampus

## 1. Introduction

Pregnancy is a vital period for offspring brain development and the trajectory of this growth may be impacted by disruptions in maternal health. Human epidemiological research has revealed an association between infection during gestation and risk for neurodevelopmental disorders (Babulas et al., 2006; Estes and McAllister, 2016; Sørensen et al., 2008). With the outbreak and worldwide spread of coronavirus disease 2019 (COVID-19), the long-lasting risks that gestational infections pose to offspring has gained significant attention (Cavalcante et al., 2020). It is critical to understand the underlying pathogenic mechanisms associated with these infections, and potential therapeutic interventions.

In preclinical research, rodent models of maternal immune activation (MIA) have been widely used to investigate biological mechanisms that underlie behavioral and cognitive abnormalities relevant to psychopathology (Estes and McAllister, 2016; Kentner et al., 2019a; Knuesel et al., 2014; Bauman & Van de Water, 2020). For example, a mid-gestational injection of the viral mimetic polyinosinic:polycytidylic acid (poly (I:C)), a commercially available synthetic analog of double-stranded RNA, can induce an extensive collection of innate immune responses (Mueller et al., 2019) and lead to a wide array of abnormalities in brain morphology (Li et al., 2009; Meyer et al., 2006) as well as neurochemical and pharmacological reactions (Zuckerman et al., 2003; Zuckerman and Weiner, 2005) associated with altered behaviors and cognitive abilities (Li et al., 2014; Ozawa et al., 2006). Despite the abovementioned progress in unravelling the disruptive effects of MIA, we still have limited understanding of how gestational immune insults alter the neurobiological substrates that underlie the behavioral abnormalities relevant to psychopathology, and more importantly, the potential for protective strategies against MIA-induced neurodevelopmental abnormalities.

At the neural level, the ventral hippocampus is a potentially important region for mediating MIA’s disruptive effects on social interaction, anxiety, and physiological responses to stress. The ventral hippocampus is an anatomically heterogeneous brain region that demonstrates remarkable plasticity (Fanselow and Dong, 2010). This region has been implicated in modulating social interactions (Bagot et al., 2015; Felix-Ortiz and Tye, 2014), anxiety-like behaviors, and regulation of the stress response (Gulyaeva, 2019; McEwen et al., 2016). In anxiogenic environments, increased neural activity of the ventral, but not dorsal, hippocampus is associated with elevated displays of anxiety-related behaviors (Adhikari et al., 2010, 2011). This association is mediated, at least partially, by synchronized neural activity between the ventral hippocampus and the prefrontal cortex (PFC), to promote anxiety-related behaviors (Adhikari et al., 2010, 2011; Padilla-Coreano et al., 2016). Previous work has demonstrated disruptive effects of MIA on several aspects of hippocampal anatomy and functioning, including decreased levels of glucocorticoid receptors and glutamate (Connors et al., 2014), altered glutamate decarboxylase expression (Dickerson et al., 2014) and lower glucose uptake (Hadar et al., 2015). Together, this work implicates the ventral hippocampus as an important brain site for mediating MIA’s disruptive effects on social behaviors and stress responses.

One characteristic of MIA’s effects is that many phenotypic abnormalities are not fully developed until late in adolescence or early adulthood (Ozawa et al., 2006; Piontkewitz et al., 2011; Zuckerman et al., 2003), which gives a time window for the application of potential therapeutic interventions. However, there has been relatively little evaluation on how supportive measures may protect the brain and behavior (Kentner et al., 2019b; Luby et al., 2020). Environmental enrichment (EE) is a non-invasive and non-pharmacological therapy characterized by exposure to novel environments with rich social, motor, cognitive and sensory stimulation. Growing evidence has shown that exposure to an enriched environment can enhance brain plasticity (e.g., increased dendritic branching, synaptogenesis, neurogenesis etc; Brenes et al., 2016; Kolb et al., 1998; Van Praag et al., 2000), the turnover of several neurotransmitters (Escorihuela et al., 1995; Hilario et al., 2016), as well as improvement in cognitive functions (Williams et al., 2001; Zeleznikow-Johnston et al., 2017). In humans, EE has been used to reverse behavioral and cognitive impairments associated with stroke, cerebral palsy, and autism (Aronoff et al., 2016; de Brito Brandã et al., 2019; Janssen et al., 2014; Morgan et al., 2014; Morgan et al., 2015; Rosbergen et al., 2017; Woo et al., 2015). Using animal models of health and disease to explore these benefits of EE provide insight into the underlying biological mechanisms of its successes and limitations.

In the animal laboratory, Van Dellen et al., (2000) were the first to demonstrate that living in spatially complex environments (i.e., EE) delayed the appearance of neurological symptoms of Huntington’s disease, and thereafter EE has been shown to improve brain pathology in animal models of Alzheimer’s disease and major depression (Chourbaji et al., 2011; Herring et al., 2009). A previous study using a poly (I:C) mouse model found that MIA interfered with the supportive effects of post-weaning EE (Buschert et al., 2016). However, this study employed male CD-1 mice that become aggressive when housed in EE (McQuaid et al., 2012; McQuaid et al., 2013; McQuaid et al., 2018). The experience of unstable social hierarchies’ and territorial aggression (e.g., fighting for resources/EE devices) likely created a negative, as opposed to enriching, environment.

Our lab has previously used rat models of prenatal lipopolysaccharide (LPS) administration to assess the beneficial effects of EE (Connors et al., 2014; Kentner et al., 2016; Kentner et al., 2018a; Núñez Estevez et al., 2020; Zhao et al., 2020) and its beneficial effects may be mediated through its interaction with maternal care, which impacts offspring development via nursing behaviors and nutritional provisioning, in addition to temperature regulation (Francis and Meaney, 1999; Connors et al., 2015). Although previous studies on the protective effects of EE revealed ambiguous findings regarding maternal nursing behaviors (e.g., licking, grooming, and crouching; Begenisic et al., 2015; Cutuli et al., 2015; de Jong et al., 1998; Li et al., 2016; Sale et al., 2004; Strzelewicz et al., 2019; Welberg et al., 2006; Zuena et al., 2016), enriched dams generally showed better nest-building quality (Cutuli et al., 2015). In contrast to EE’s positive effects, gestational treatment with poly (I:C) led to deficient maternal care behaviors (Ronovsky et al., 2017; Schwendener et al., 2009). Collectively, prior research lends some support to the idea that EE may interact with parental care to offset effects of early health adversities by buffering hypothalamic pituitary adrenal (HPA) axis dysregulation. Indeed, when used in pediatric clinical settings, the benefits of EE are mediated at least in part by caregiver engagement (Aronoff et al., 2016; Bowman & Evans, 2019; de Brito Brandão et al., 2019; Morgan et al., 2014; Morgan et al., 2015; Woo et al., 2015).

Building upon our previous work in rats, the current study aimed to examine if the protective effects of EE on social behavior and HPA axis regulation extends to toll-like 3 receptor activation by using a poly (I:C)-induced MIA model in mice. Importantly, we used C57BL/6J mice, which are a less aggressive strain; they did not demonstrate evidence of increased fighting in response to higher cage densities and competition for resources (Nicholson et al., 2009), suggesting that EE is appropriate for these animals.

## 2. Materials and methods

### 2.1. Animals and housing

C57BL/6J mice were acquired from the Jackson Laboratory (Bar Harbor, ME), and housed at 20°C on a 12 h light/dark cycle (0700–1900 light) with ad libitum access to food and water. A schematic timeline of experimental procedures is presented in **Figure. 1A**. Female mice were housed in pairs in one of two conditions: environmental enrichment (EE; N40HT mouse cage, Ancare, Bellmore, NY; see **Figure. 1B**), comprised of a larger cage and access to toys, tubes, a Nylabone, Nestlets^®^ (Ancare, Bellmore, NY) and Bed-r’ Nest^®^ (ScottPharma Solutions, Marlborough MA), or standard cages (SD; N10HT mouse cage, Ancare, Bellmore, NY; see **Figure. 1C**) with Nestlets^®^ and a Nylabone only. Male animals were paired in SD conditions unless they were breeding, at which point they were housed with two females in EE or SD cages. Immediately after breeding, dams were placed into clean cages, maintaining their assigned housing conditions.

**Figure 1.**
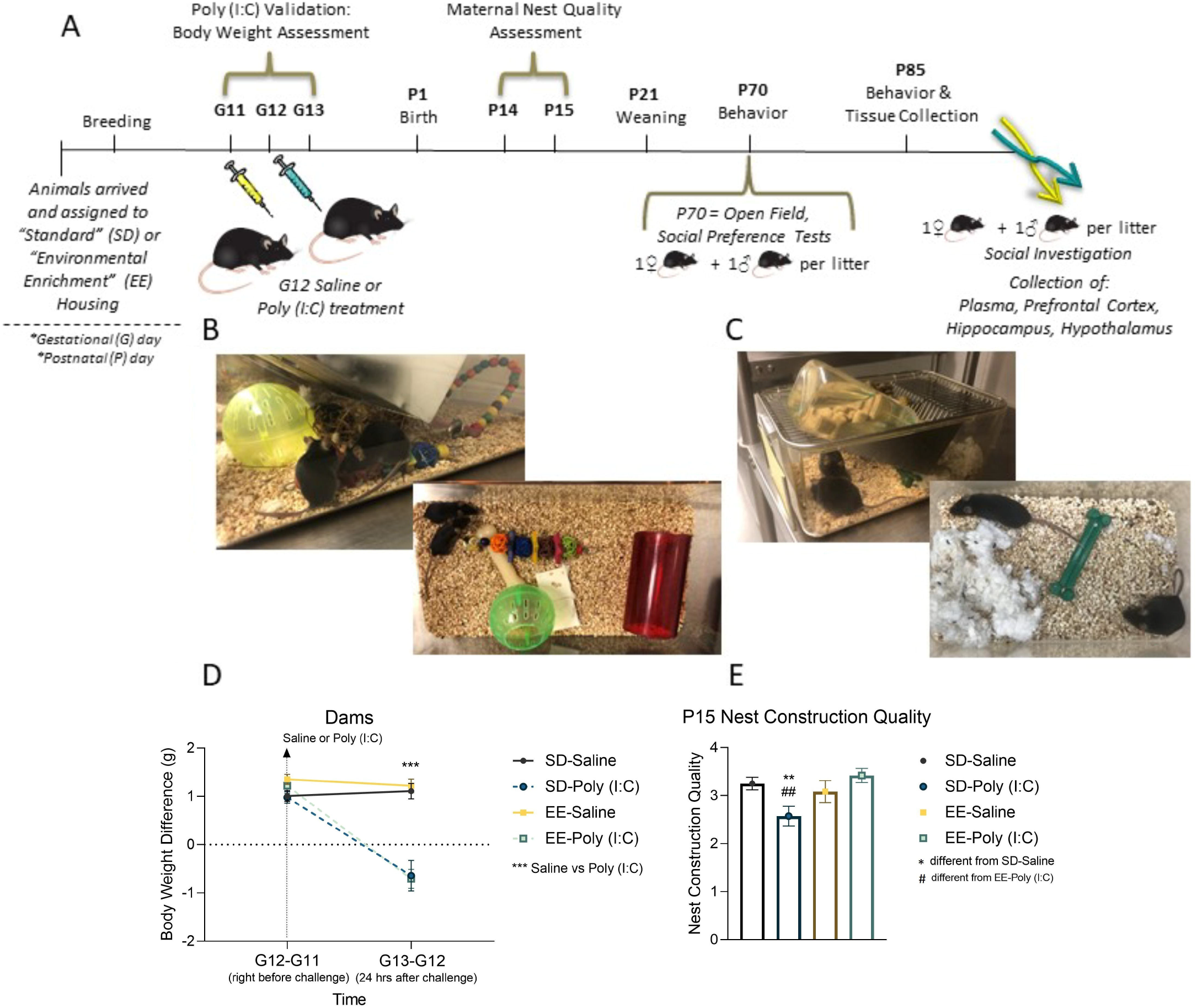
Experimental Timeline and Maternal Measures. (**A)** Flow chart of experiment procedures. Pictures of the **(B)** environmental enrichment and **(C)** standard housing conditions. **(D)** Maternal body weight differences (grams) between gestational days (G) 11 to 12 and days 12 to 13, and **(E)** postnatal day 15 nest quality scores following maternal immune activation (MIA) and environmental enrichment housing. Data are expressed as mean ± SEM, n=12-14 dams per MIA and housing group. *p < 0.05, **p < 0.01, ***p <0.001, versus SD-saline; #p < 0.05, ##p < 0.01, ###p <0.001, versus EE-poly (I:C).

On the morning of gestational day (G)12, pregnant dams were randomly allocated to receive a 20 mg/kg intraperitoneal injection of polyinosine-polycytidylic acid (poly (I:C); tlrl-picw, lot number PIW-41-03, InvivoGen), or vehicle (sterile pyrogen-free 0.9% NaCl). Twenty-four hours later, maternal body weights were recorded to validate the poly (I:C) challenge by either slowed weight gain or loss. To ensure the safety of the pups, toys and Nylabones^®^ were removed on G19 and returned after maternal nest quality evaluations on P15. To maintain standardized litter sizes, some pups were fostered to same age and group matched dams. Additional methodological details can be found in the reporting table from Kentner et al. (2019a), provided as **Supplementary Table S1**. Animal procedures were approved by the MCPHS University Institutional Animal Care and Use Committee and carried out in compliance with the recommendations outlined by the Guide for the Care and Use of Laboratory Animals of the National Institutes of Health.

## 2.2. Nest building quality

While there are some inconsistencies in the literature, some investigators find that gestational poly (I:C) alters parental care quality (e.g., licking/grooming, nursing, time off nest) (Weber-Stadlbauer et al., 2020; Ronovsky et al., 2017; Schwendener et al., 2009). Enriched environments are also known to change patterns of maternal behavior in rats, and in some cases seem to make dams more efficient in their care duties (Baldini et al, 2013; Connors et al., 2015; Strzelewicz et al., 2019; Welberg et al., 2006). Extensive maternal care observations are difficult with our mouse model as the dams tend to build closed dome-shaped nests, hiding their pups and obscuring maternal-pup interactions. Therefore, we conducted passive maternal nest quality observations on P15. This was done 24 hours after litters were placed into a fresh cage and each dam provided four fresh Nestlets^®^ (Ancare, Bellmore, NY). Nest quality was scored on a 5-point Likert scale, adapted from Gerfen et al., (2006) in which 0 = no nest; 1 = flat nest; 2 = a slightly cupped shape but less than 1/2 of a dome shape; 3 = taller than 1/2 of a dome shape, but not fully enclosed and 4 = fully enclosed nest/dome. We also recorded the number of unused Nestlets^®^.

### 2.3. Offspring behavior

Offspring were weaned into same-sex groups on P21 and maintained in their housing assignments until behavioral tests commenced (SD = 2-3 animals/cage; EE = 4-5 animals/cage). **Supplementary Table S2** outlines the litters and treatment groups for the study. On P70, one male and one female from each litter was habituated to an open field arena (40 cm×40 cm×28 cm) for three-minutes (Duque-Wilckens et al., 2020; Williams et al., 2020). Behavior was recorded and videos scored via an automated behavioral monitoring software program (Cleversys TopScan, Reston, VA) for percent time spent in the center of the arena and total distance traveled (mm). Rearing and grooming behaviors were evaluated using the same procedures as the social tests described below. All equipment was thoroughly cleaned with Quatriside TB between each animal and test. Immediately after the open field habituation period, two cleaned wire containment cups were placed on opposite ends of the arena for the five-minute social preference test. One cup contained a novel untreated adult mouse of the same sex, age, and strain and the other cup contained a novel object. Placement of novel mice and objects was interchanged between trials. A mouse was actively investigating when its nose was directed within 2 cm of a containment cup or it was touching the cup. For each animal, a social preference index was calculated by the formula ([time spent with the mouse] / [time spent with the inanimate object + time spent with the mouse])□-0.5 (Scarborough et al., 2020).

To determine neuronal activation associated with a social experience, animals were re-introduced to the open field arena on P85, but this time only one wire cup containing an unfamiliar mouse of the same sex, age, and strain was presented. Duration of time animals spent actively investigating the novel mouse was evaluated across the 10-minute social exposure period. We also assessed time spent in the interaction zone and vigilance behavior during the first three-minutes of the social exposure (adapted from Duque-Wilckens et al., 2020; Williams et al., 2020). Briefly, social interaction was defined as the time animals spent within 8 cm (interaction zone) of the novel mouse. A mouse was demonstrating social vigilance when its head was oriented towards the novel mouse when outside of the interaction zone. Blinded observers evaluated all social behavior videos for the frequency of rearing behavior and duration of grooming and social behaviors using a manual behavioral scoring software program (ODLog™ 2.0, http://www.macropodsoftware.com/). Inter-rater reliability was determined by Pearson *r* correlation to be 0.860–0.900 for each manually scored behavior.

### 2.4. Tissue collection and analysis

Ninety minutes after the P85 social investigation exposure, brains were collected for *c-Fos* immunohistochemistry and qPCR analyses. A mixture of Ketamine/Xylazine (150 mg/kg, i.p/15 mg/kg, i.p) was used to anesthetize animals. Blood was collected via cardiac puncture and placed into an ethylenediaminetetraacetic acid (EDTA)-coated microtainer tube (Becton Dickson, Franklin Lakes New Jersey). Samples were centrifuged at 1000 relative centrifugal force (rcf) for 10 min for plasma separation. Animals were perfused intracardially with a chilled phosphate buffer solution. Prefrontal cortex, hippocampus, and hypothalamus were dissected from the left hemisphere. Samples were frozen on dry ice and stored at −75°C until processing. The right hemisphere was post-fixed in a 4% paraformaldehyde, phosphate buffer 0.1M solution (BM-698, Boston BioProducts) overnight at 4°C. Tissue was then submerged in ice cold 10% sucrose in PBS (with 0.1% sodium azide) and incubated at 4°C overnight. The following day, solution was replaced with 30% sucrose in PBS for 3 days. Tissue was rapidly flash frozen with 2-methylbutane (O3551-4, Fisher Scientific) and stored at −75°C until sectioning.

### 2.5. Corticosterone assay

Plasma samples were evaluated with a corticosterone ELISA kit (#ADI-900-097, Enzo Life Sciences, Farmingdale, NY). The small sample assay protocol was followed, as recommended by the manufacturer, and each sample was processed in duplicate. The minimum detectable concentration was 26.99 pg/ml, and the intra- and inter-assay coefficients of variation were 6.6% and 7.8%, respectively.

### 2.6. RT-PCR

Total RNA was extracted from frozen tissue using the RNeasy Lipid Tissue Mini Kit (Qiagen, 74804) and resuspended in RNase-free water. Isolated RNA was quantified utilizing a NanoDrop 2000 spectrophotometer (ThermoFisher Scientific). According to the manufacturer’s protocol, total RNA was reverse transcribed to cDNA with the Transcriptor High Fidelity cDNA Synthesis Kit (#5081963001, Millipore Sigma) and the final cDNA solution was stored at -20°C for analysis. Quantitative real-time PCR with Taqman™ Fast Advanced Master Mix (#4444963, Applied Biosystems) was used to measure the mRNA expression of corticotropin releasing hormone (Crh, Mm01293920_s1), Crh receptor 1 (Crhr1, Mm00432670_m1), glucocorticoid receptor (Nr3c1, Mm00433832_m1), oxytocin (Oxt, Mm00726655_s1), oxytocin receptor (Oxytr, Mm01182684_m1), vasopressin receptor (Avpr1a, Mm00444092_m1x), calcium/calmodulin dependent protein kinase II alpha (Camk2a, Mm00437967_m1), and protein kinase C alpha (Prkca, Mm00440858_m1). All reactions were analyzed in triplicate using optical 96-well plates (Applied Biosystems StepOnePlus™ Real-Time PCR System) and relative gene expression levels were evaluated using the 2 ^−^ΔΔ^CT^ method with 18S (Hs99999901_s1). This housekeeping gene was selected as it was not affected by MIA or housing condition. Gene expression was normalized in relation to 18S and data presented as mean expression relative to same sex SD-saline treated controls.

### 2.7. Immunohistochemistry

Coronal sections (40 um thick) were obtained on a cryostat (Leica CM1860) as a 1:4 series and stored at -20°C in cryoprotectant until immunocytochemistry. Free-floating sections were washed in PBS to remove the cryoprotectant prior to incubation in rabbit anti-cFos primary antibody (1:5000; ABE457, Millipore Sigma) for one hour at room temperature (RT) and 48 hours at 4°C. Tissue was then washed in PBS and incubated in goat anti-rabbit IgG biotinylated secondary antibody (1:600) for one hour at RT. Tissue was washed in PBS and incubated in A/B solution (A and B solutions of the Vectastain® Elite ABC-HRP Kit, #PK-6161, Vector Laboratories) for one hour at RT. This was followed by another PBS wash after which tissue was briefly washed in 0.175 M sodium acetate and incubated with 3,3′-Diaminobenzidine tetrahydrochloride (Ni-DAB; #D5905-50TAB, Millipore Sigma). Sections were then washed briefly with 0.05 M sodium acetate and PBS and stored at 4°C in PBS until mounting. Tissue was mounted in 0.85% saline and slides air-dried for 48 hours before being dehydrated in an alcohol dilution series, cleared in xylene, and cover slipped with DPX (#06522-100ML, Millipore Sigma). Slides were imaged on a Keyence Microscope [BZ-X800, Keyence] and the number of Fos-immunoreactive cells in each region of interest was counted by two experimenters blind to experimental treatments during counting. Inter-rater reliability was determined by Pearson *r* correlation to be 0.830–0.990 for each brain region evaluated. The area of each selected region was measured with NIH ImageJ software (Schneider et al., 2012) to determine the number of stained cells per square millimeter. Prelimbic and infralimbic regions of the medial prefrontal cortex were identified using the Allen Mouse Brain Atlas (image 36), as were regions of the ventral hippocampus (e.g., vCA1, CA2, vCA3, dentate gyrus), the supramammillary nucleus (SuM) and whole hypothalamus (image 81).

### 2.8. Statistical analysis

Statistics were performed using the software package Statistical Software for the Social Sciences (SPSS) version 26.0 (IBM, Armonk, NY). The assumption of normality was evaluated with the Shapiro-Wilk test and Kruskal-Wallis tests (expressed as *X^2^*) employed in rare cases of significantly skewed data. Maternal body weights were evaluated using repeated measures 2-way ANOVA (MIA × housing). Since qPCR data were normalized to same sex controls, a 2-way ANOVA (MIA × housing) was conducted for each gene of interest in each sex separately. The remaining measures were evaluated with 3-way ANOVAs (sex × MIA × housing). Pearson correlations were analyzed between the P85 social measures versus ventral hippocampal Fos -immunoreactivity and gene expression. LSD post hocs were applied except where there were fewer than three levels, in which case pairwise t-tests and Levene’s (applied in the occurrence of unequal variances) were utilized. The False Discovery Rate (FDR) was applied to correct for multiple testing procedures in all gene expression and correlation experiments. All data are graphically expressed as mean ± SEM. If there were no significant sex differences, data were collapsed together for visualization. The partial eta-squared (*n*_*p*_^2^) is also reported as an index of effect size for the ANOVAs (the range of values being 0.02 = small effect, 0.13 = moderate effect, 0.26 = large effect; Miles and Shevlin, 2001).

## 3. Results

### 3.1. MIA challenge interferes with maternal body weight gain and nesting behaviors while EE supports maternal care quality

To validate the integrity of our MIA model, we evaluated maternal body weight gain following gestational poly (I:C) challenge. While baseline weight gain was comparable across conditions (p>0.05), dams treated with poly (I:C) had considerably slower body weight gain compared to saline (p=0.001; **Figure 1D**) 24 hours after challenge (*time x MIA*: F(1, 46) = 27.433, p = 0.001, *n*_*p*_^*2*^ = 0.374). This provides evidence that our poly (I:C) administration protocol induced an immune response (Kentner et al., 2019a).

On P14, dams were given four Nestlets® and nest construction quality evaluated 24 hours later. There were no differences across treatments in terms of the number of nesting resources used (p>0.05; data not shown). However, SD-poly (I:C) dams constructed nests of significantly poorer quality compared to SD-saline (*X*^*2*^(1) = 6.378, p = 0.012) and EE-poly (I:C) mice (*X*^*2*^(1) = 7.84, p =0.005; **Figure 1E**). There were no differences between EE-saline and EE-poly (I:C) nest quality (p>0.05), highlighting the protective effects of enhanced environments on parental care.

### 3.2. Environmental enrichment blocks prenatal poly (I:C) induced reductions in social motivation and associated displays of repetitive behavior

While there is a strong focus on identifying the negative consequences and mechanisms of MIA on the developing brain and behavior, very little attention is directed towards the prevention or rehabilitation of these effects in laboratory animal studies (Kentner et al., 2019b). Given EE’s protective effects in the LPS-indued MIA model (Nunez et al., 2020; Connors et al., 2014), we determined whether this protection extended to the poly (I:C) model of MIA. As expected, P70 social preference was significantly reduced by poly (I:C)-induced MIA in both SD males (**Figure 2A**) and females (**Figure 2B**; *MIA x housing*: F(1, 83) = 7.047, p = 0.010, *n*_*p*_^*2*^=0.078). This effect was completely blocked by enriched housing (SD-saline vs SD-poly (I:C): p = 0.005; SD-poly (I:C) vs EE-poly (I:C): p = 0.0001; **Figure 2A-C**).

**Figure 2.**
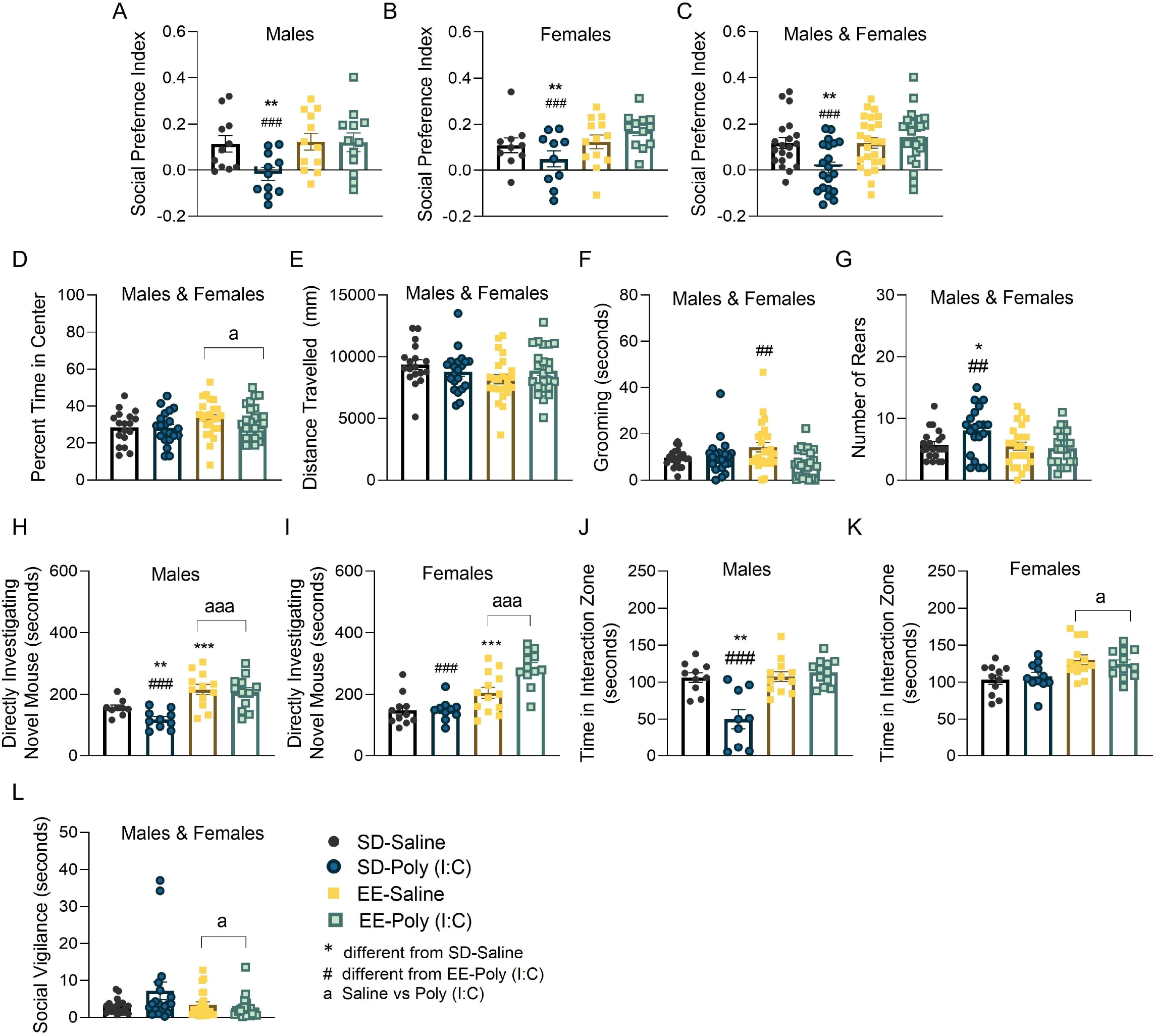
The effects of maternal immune activation (MIA) and environmental enrichment on adult offspring social behavior. **(A-G)** Represent data from the postnatal day (P) 70 social preference test. Social preference index for **(A)** male, **(B)** female and **(C)** both sexes combined. **(D)** Percent of time in the center and **(E)** distance traveled (mm) in the open field test for male and female offspring. **(F)** The duration (seconds) of time spent grooming and **(G)** the number of rears. **(H-L)** Represent data from the P85 social investigation test. The duration of time (seconds) spent directly investigating a novel conspecific for **(H)** male and **(I)** female offspring. Time that **(J)** males and **(K)** females spent in the interaction zone during the social investigation experience. **(L)** Time that male and female mice spent in social vigilance. Data are expressed as mean ± SEM, n = 9-13 litters represented per sex, MIA, and housing group. *p < 0.05, **p < 0.01, ***p <0.001, versus SD-saline; #p < 0.05, ##p < 0.01, ###p <0.001, versus EE-poly (I:C); ^a^p < 0.05, ^aa^p < 0.01, ^aaa^p < 0.001, main effect of housing.

To determine whether the effect of MIA on social interest was reflective of an increased anxiety-like phenotype, we evaluated behaviors in the P70 open field test. During this assessment it was apparent that only housing condition (F(1,83) = 4.024, p =0.048, *n*_*p*_^*2*^ = 0.049) impacted the percentage of time spent in the center of the arena (EE: 32.69±1.36 vs SD 29±1.39; p = 0.028; **Figure 2D**). Distance traveled was not affected by any treatment condition (p>0.05; **Figure 2E**). Male animals reared more frequently than females (p<0.05; **Supplementary Figures S1A,B**; *P70 main effect of sex:* F(1, 83) = 5.518, p = 0.021, *n*_*p*_^*2*^ = 0.062; *P85 main effect of sex:* F(1,81) = 4.436, p =0.038, n_p_^2^ = 0.051) and female EE mice groomed more than SD females (p = 0.0001; **Supplementary Figures S1C,D**; *sex x housing:* F(1,81) = 4.352, p =0.040, *n*_*p*_^*2*^ = 0.050) in the open field test, but there was no effect of MIA on these measures. EE animals tend to display higher levels of body licking behavior as they habituate more quickly to novel environments (Rojas-Carvajal et al., 2018). Here, EE-saline mice groomed longer during the social preference test, an effect that was blunted by MIA (*EE-saline vs EE-poly (I:C): X*^2^(1) = 7.899, p = 0.005; **Figure 2F**). However, there were no group differences in grooming during the P85 social investigation test (p>0.05; **Supplementary Figure S1E**).

In contrast, during the P70 social preference test, male and female SD-poly (I:C) mice showed a higher frequency of rearing behaviors compared to same-sex SD-saline (p = 0.024) and EE-poly (I:C) mice (p = 0.006; **Figure 2G**; *MIA x housing*: F(1, 83) = 4.313, p = 0.035, p = 0.035, *n* _*p*_ ^*2*^ = 0.054).

To more fully characterize the heightened stress response/delayed stress recovery associated with MIA, we exposed animals to a social stimulus on P85 to correlate behavior with *c-Fos* activation and gene expression ninety minutes later. Male SD-poly (I:C) mice spent less time directly engaging with the novel mouse compared to SD-saline males (p = 0.010) across the 10 minute test period (*sex x MIA*: (F(1, 81) = 9.209, p = 0.003, *n*_*p*_^*2*^=0.102; *MIA x housing*: F(1, 81) = 5.546, p = 0.012, *n*_*p*_^*2*^ = 0.075; **Figure 2H,I**). As expected, male and female EE animals spent more time in direct social contact compared to SD mice (p = 0.001), blocking the attenuating effect of poly (I:C) on social behavior. Patterns of poly (I:C) impeding male social interest (p = 0.002), and EE increasing social interest (p = 0.0001), were again confirmed by the respective reduced and elevated times spent within the interaction zone (*sex x MIA x housing:* F(1, 81) = 11.162, p = 0.001, *n*_*p*_^*2*^ = 0.120; **Figure 2J,K**). While social vigilance was not directly impacted by MIA (p>0.05), it was significantly reduced by EE housing (*SD:* 4.96±1.16 *vs EE:* 3.10±0.46; *X*^*2*^(1) = 4.171, p = 0.041; **Figure 2L**), which may account for higher instances of social contact from these animals.

### 3.3. MIA reduced c-Fos activation in the dentate gyrus of male and female offspring

Ninety minutes following the P85 social exposure, we evaluated *c-Fos* activation throughout brain regions critical to anxiety-like and social behavior (Ko, 2017; Chen et al., 2020; Walsh et al., 2020; **Figure 3A,B**). There were no main effects or interactions with respect to sex, so male and female data were collapsed and visualized together. Although *c-Fos* activation in the medial prefrontal cortex (prelimbic and infralimbic regions; **Figure 3C)** and hippocampal CA1, CA2, and CA3 regions were not affected (p > 0.05; **Figure 3D**), in the hippocampus MIA significantly decreased the number of *c-Fos* immunoreactive cells expressed in the dentate gyrus (DG; *X*^*2*^(1) = 4.001, p = 0.0.045; *MIA*: x□ = 97.35±12.68 *vs Saline*: x□ = 174.91±31.98; **Figure 3D**). The DG is important for the mediation of odor and reward processing, in addition to social recognition and social avoidance (Kesner et al., 2018; Kheirbek et al., 2013; Weeden et a., 2015).

**Figure 3.**
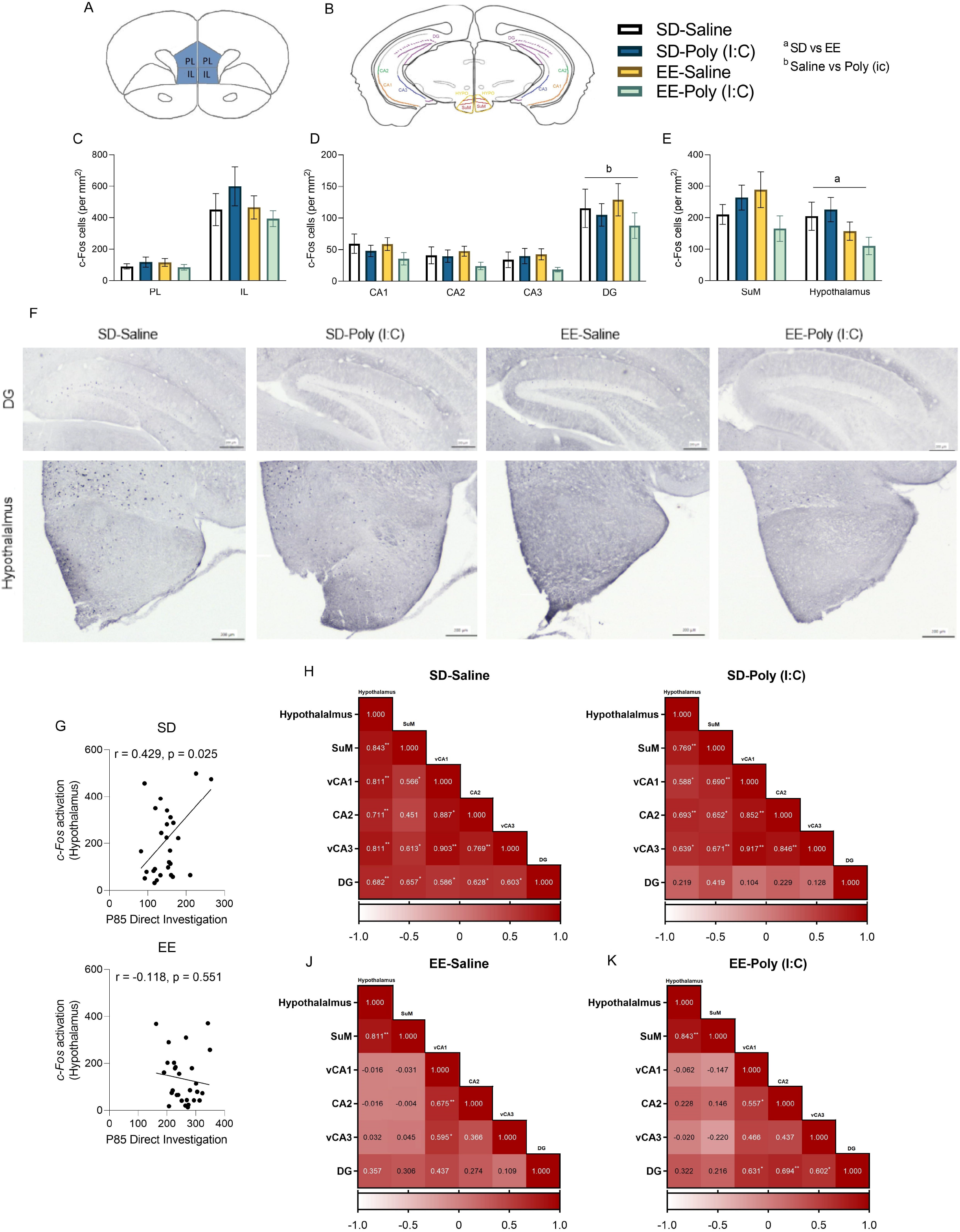
Central *c-Fos* activation following a postnatal day (P) 85 social exposure in male and female offspring exposed to maternal immune activation (MIA) and environmental enrichment (EE). **(A)** Coronal section of the medial prefrontal cortex, including prelimbic (PL) and infralimbic (IL) regions. **(B)** Coronal section of the hippocampus and hypothalamus including CA1, CA2, CA3, dentate gyrus (DG), the supramammillary nucleus (SuM) and whole hypothalamus. Mean *c-Fos* (per mm^2^) in the **(C)** medial prefrontal cortex PL and IL regions, **(D)** hippocampus (CA1, CA2, CA3, DG), and **(E)** SuM and whole hypothalamus following a ten-minute exposure to a novel social conspecific. **(F)** Representative images of *c-Fos* immunostaining in the dentate gyrus (top) and hypothalamus, including SuM (bottom) of standard housed (SD)-saline, SD-poly (I:C), EE-saline and EE-poly (I:C) male and female offspring. Scale bar equals 200 μm. **(G)** Pearson correlations of *c-Fos* activation and time (seconds) spent in direct social investigation, for SD (top) and EE (bottom) mice. Correlated neural activity patterns computed from Pearson’s bivariate correlations of *c-Fos* positive cells between the hypothalamus (including SuM) and hippocampal regions for **(H)** SD-saline, **(I)** SD-poly (I:C), **(J)** EE-saline, and **(K)** EE-poly (I:C) mice. Mean ± SEM. There were no significant sex differences so male and female data were collapsed together; n=13-17 total per group. ^a^p < 0.05, ^aa^p < 0.01, main effect of housing; ^b^p < 0.05, ^bb^p < 0.01, main effect of MIA; *p <0.05, **p<0.01, significant Pearson’s correlations

### 3.4. Environmental enrichment housing remodels neuronal activation patterns associated with a novel social experience

Neurons in the supramammillary nucleus (SuM) are activated during novel social interactions (Chen et al., 2020). Specifically, contextual novelty reportedly activates the SuM-dentate gyrus circuit and social novelty activates a SuM-CA2 circuit (Chen et al., 2020; Walsh et al., 2020). This led us to evaluate *c-Fos* immunoreactivity in this hypothalamic region following MIA. While evaluation of the SuM revealed a significant MIA by housing interaction (F(1, 50) = 4.098, p = 0.048, *n*_*p*_^*2*^ = 0.076; **Figure 3E**), follow-up tests were not significant. However, housing significantly affected *c-Fos* activation in the hypothalamus (including the SuM region; *X*^*2*^(1) = 4.137, p = 0.042; **Figure 3E**) during the P85 social exposure. The lower neural activation of EE animals (x□ = 133.87±20.25) in this region may be indicative of their faster habituation rates compared to SD (x□ = 214.23±29.64; Rojas-Carvajal et al., 2018). **Figure 3F** shows representative samples of *c-Fos* staining for each experimental group.

Pearson’s correlations confirmed that direct social investigation of the novel mouse on P85 was positively associated with elevated *c-Fos* immunoreactivity in the hypothalamus of standard housed animals (r = 0.429, p = 0.025; **Figure 3G**); this effect was blunted in EE mice (p >0.05; **Figure 3G**).

Hypothalamic *c-Fos* activity was not correlated with time spent in the interaction zone, or social vigilance (p>0.05), highlighting the specificity of this activation. During the P85 social exposure, neural activation in the hypothalamus (including the SuM) was strongly and positively correlated with vCA1 (r = 0.734, p = 0.0001), CA2 (r = 0.703, p = 0.0001), vCA3 (r = 0.736, p = 0.0001), and DG (r = 0.725, p = 0.0001) in SD animals. However, these putative circuit associations were lost with a more complex EE housing condition (p >0.05). *A priori* tests of MIA and housing interactions among these associations again demonstrated strong positive correlations between hypothalamus + SuM with regions vCA1, CA2, vCA3 and DG in SD-saline housed mice (p <0. 05; **Figure 3H)** during the P85 social exposure. Most of these associations were maintained with MIA treatment in SD animals (**Figure 3I**), except for the relationship with the DG which was lost (p >0.05). Interestingly, EE housing disrupted many of the associations observed in SD animals (**Figure 3J,K)**. This may be an important consideration when trying to understand complex neural circuits using simplistic environmental conditions (Kentner et al., 2018c).

### 3.5. MIA-induced dysregulation of stress associated markers in the ventral hippocampus is antagonized by environmental enrichment

Following the P85 social exposure, several mRNA expression patterns emerged throughout the brain as a function of MIA. While there was no treatment effect on hippocampal expression of Oxt mRNA (p>0.05; **Supplementary Table S3**), Oxtr was elevated in SD-poly (I:C) males (p = 0.003) and lowered in SD-poly (I:C) females (p = 0.029) compared to their respective same-sex SD-saline and poly (I:C) enriched counterparts (*males:* (F(1, 28) = 6.383, p = 0.017, *n*_*p*_^*2*^ = 0.186); *females*: (F(1, 28) = 7.912 p = 0.009, *n*_*p*_^*2*^ = 0.220**; Figure 4A,B**).

**Figure 4.**
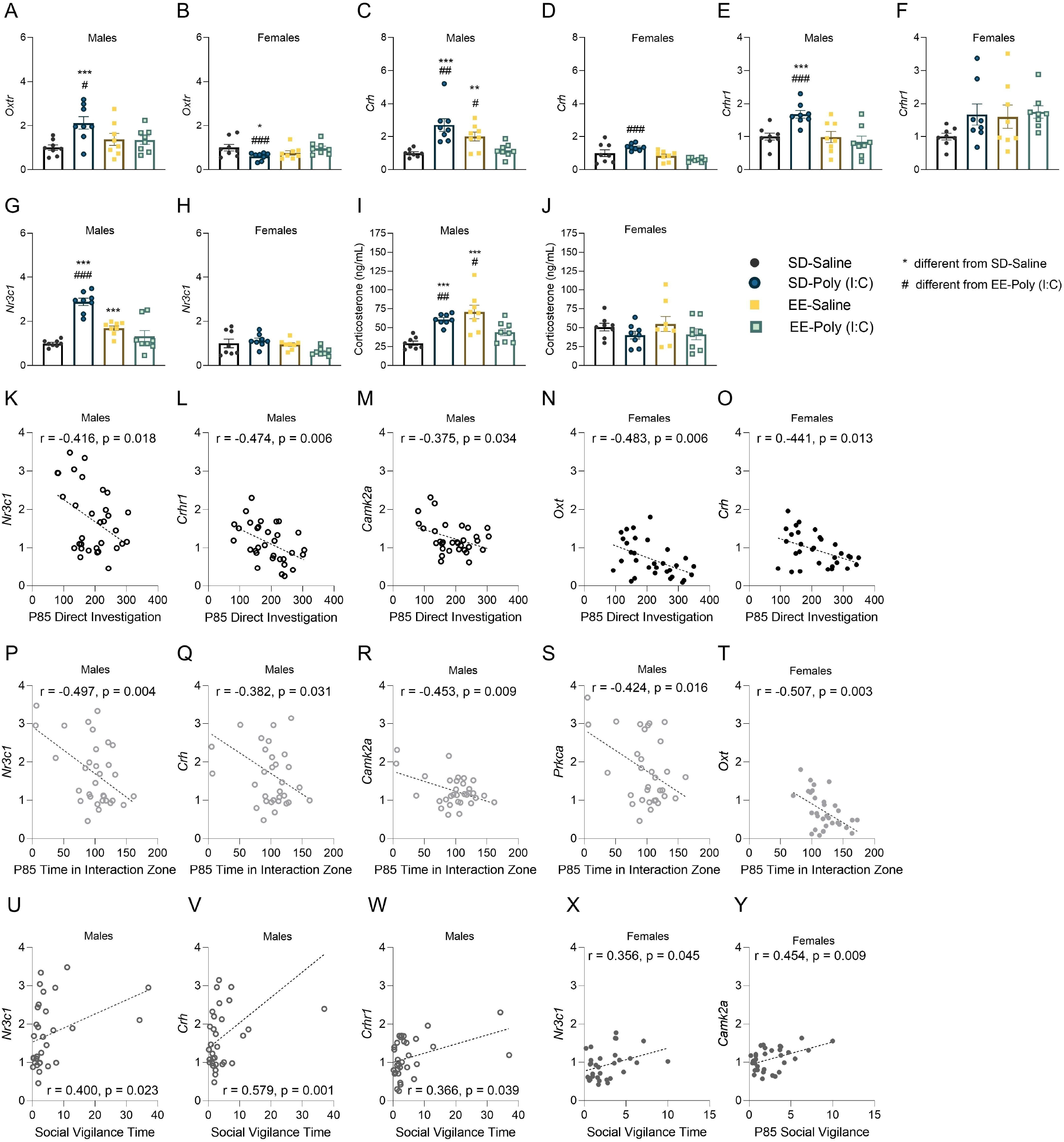
Ventral hippocampal gene expression following maternal immune activation (MIA) and environmental enrichment housing. Male and female offspring levels of **(A,B)** Oxtr, **(C,D)** Crh, **(E, F)** Crhr1, **(G, H)** and Nr3c1 expressed as relative mRNA expression and **(I, J)** plasma corticosterone. Pearson correlations of ventral hippocampal relative mRNA expression and time (seconds) spent in **(K-O)** direct social investigation, **(P-T)** the interaction zone, and **(U-Y)** social vigilance on postnatal day 85. Gene expression data are expressed as Mean ± SEM, n=8 litters represented per sex, MIA, and housing group. *p < 0.05, **p < 0.01, ***p <0.001, versus SD-saline; #p < 0.05, ##p < 0.01, ###p <0.001, versus EE-poly (I:C).

Male (F(1, 28) = 25.143, p = 0.001, *n*_*p*_^*2*^ = 0.473; **Figure 4C**) and female hippocampal Crh (F(1, 28) = 5.673, p = 0.024, *n*_*p*_^*2*^ = 0.168; **Figure 4D**) was also associated with significant MIA by housing effects. Specifically, SD-poly (I:C) mice had elevated levels of Crh compared to SD-saline (*males*: p=0.001) and EE-poly (I:C) animals (*males*: p = 0.003; *females*: p= 0.0001). Male EE-saline mice had higher Crh compared to SD-saline (p = 0.003) and EE-poly (I:C) animals (p = 0.016) as well. Notably, SD-poly (I:C) males also had enhanced expression of Crhr1 in the ventral hippocampus (*SD-saline*: p = 0.001; *EE-poly (I:C)*: p = 0.001), while EE-saline animals did not (p>0.05; *MIA x housing*: F(1, 28) = 8.512, p = 0.007, *n*_*p*_^*2*^ = 0.233; **Figure 4E,F**). Similar patterns of sex-specific expression associated with MIA were observed for hippocampal Nr3c1 (*male MIA x housing*: F(1, 28) = 46.319, p = 0.0001, *n*_*p*_^*2*^ = 0.623); *females:* p > 0.05; **Figure 4G,H**), Prkca (*male MIA by housing*: F(1, 28) = 35.560, p = 0.0001, *n*_*p*_^*2*^ = 0.559; *female MIA x housing*: F(1, 28) = 7.153, p = 0.01, *n* _*p*_^*2*^ = 0.203; **Supplementary Table S3**), and Camk2a (*male MIA x housing*: F(1, 28) = 5.685, p = 0.024, *n*_*p*_^*2*^ = 0.169; *females:* p > 0.05; **Supplementary Table S3**). Overall, these data suggest that male SD-poly (I:C) animals are either demonstrating a heightened acute stress response and/or impaired ability to cope and recover from stress. Moreover, these effects can be blocked by enrichment housing. Additional sex, MIA and housing effects were revealed for the expression of genes in the prefrontal cortex, ventral hippocampus, and hypothalamus which can be found in **Supplementary Table S3**.

### 3.6. Glucocorticoid associated recovery following a social stress exposure is delayed in MIA offspring

Elevated plasma corticosterone level is one indicator of an active stress response (McEwen et al., 2015). Ninety minutes following the social stress exposure, male SD-poly (I:C) mice still demonstrated elevated levels of this steroid hormone compared to SD-saline (p = 0.0001) and EE-poly (I:C) animals (p = 0.008; F(1, 55) = 8.781, p = 0.004,, *n*_*p*_^*2*^ = 0.138; **Figure 4I,J**), indicative of delayed stress recovery in these mice. While enrichment housing was associated with lower plasma corticosterone in poly (I:C) treated males, EE-saline males had elevated levels of this steroid hormone (*SD-saline*: p = 0.001; *EE-poly*: 0.019) and hippocampal Nr3c1 (*SD-saline*: p = 0.0001; *EE-poly*: 0.0001; **Figure 4G**).

### 3.7. Ventral hippocampal gene expression is sex-dependently associated with P85 social investigation

Direct social investigation of the novel mouse was associated with sex-specific ventral hippocampal gene expression, supporting a role for these genes in our social behavior measures. Specifically, higher levels of Nr3c1, Crhr1, CamK2a (males), Oxt and Crh (females) were associated with lower social exploration levels (**Figure 4K-O**). Reduced time spent in the interaction zone was similarly associated with higher levels of genes in this region (males: Nr3c1, Crh, Crhr1, Camk2a, Prkca; females: Oxt; (**Figure 4P-T**). In contrast, heightened social vigilance was correlated with higher expression of several stress-associated gene markers in the ventral hippocampus (males: Nr3c1, Crh; females: Nr3c1, Camk2a, Oxytr; **Figure 4U-Y**).

## 4. Discussion

In the current study, we demonstrate that poly (I:C)-induced MIA imposes sex-specific disruptions on a variety of physiological and behavioral indices. Importantly, EE housing buffered many of these MIA effects, as reflected by more regulated displays of affiliative behaviors and a reduction in repetitive actions. Moreover, MIA associated elevations in plasma corticosterone, and several hippocampal mRNA markers indicative of HPA axis dysregulation, were dampened by the environmental intervention. Our results underscore the hippocampus as a region mediating MIA’s disruptive effects and as a target for interventions such as EE. We also observed MIA to impair maternal nest building quality, which was prevented by EE housing. This suggests that parental care may be an important mediator of EE’s benefits. Altogether, these data highlight the role of MIA in the behavioral expression and alteration of neural activities and markers associated with stress responses (Estes and McAllister, 2016; Knuesel et al., 2014). Moreover, our findings identify underlying mechanisms that support EE as a protective strategy against MIA-induced neurodevelopmental abnormalities.

### The Effects of MIA and EE on Maternal Nesting Behavior

Maternal immune activation not only affects offspring behavior, but the parental behavior of the dam as well. The poor nest quality displayed by our MIA mothers echoes previous observations on the consequences of gestational immune insult (Aubert et al., 1997). Quality nest building is critical for maintaining pup thermoregulation (Weber and Olsson, 2008). Deterioration of the nest impacts its internal temperature, which may affect the HPA-axis development of offspring (Jans and Woodside, 1990). MIA exposed dams also spend more time building their nest, and this trait can be transmitted to following generations (Berger et al., 2018; Ronovsky et al., 2017). Our data showing poorer nest quality adds to these findings and together suggests that poly (I:C) treated mothers may be less efficient or skilled in maternal care overall.

Supportive EE interventions can help buffer the effects of MIA on maternal nest building behavior. The mechanisms underlying the disrupted nest building performance remains unclear as nest-building is a complex behavior that requires fraying, pulling, sorting, and fluffing of nest materials (Gaskill et al., 2012). Changes in motivation or motor ability for any of these behaviors can impact nest building quality. In addition to MIA, other stressors experienced prenatally such as scrapie infection (Cunningham et al., 2003), or heat and restraint (Kinsley and Svare, 1988), have also impaired nest-building behaviors. This suggests that reductions in nest quality were precipitated by stress. Therefore, the protective effects of EE on nest building are likely mediated, at least in part, by enhancing parental stress resilience (Crofton et al., 2015; Lehmann and Herkenham, 2011). Prior to the nest quality evaluation, EE dams had been provided with additional nesting materials (i.e., Bed-r-Rest discs + Nestlets®). This supplemental support may have primed EE-poly (I:C) dams to be more responsive to the materials available to them during the nest construction test. Indeed, a previous report showed that extra naturalistic nesting materials prompted mice to construct complex dome-shaped nests, similar to those in the wild (Hess et al., 2008) and what we tended to see in our SD-saline and EE dams here.

### The Effects of MIA and EE on Offspring Social Behavior

MIA exposed mice demonstrated evidence of social aversion which was buffered by EE housing. In line with previous evidence (Choi et al., 2016; Hsiao et al., 2012; Hsiao et al., 2013; Scarborough et al., 2020; Schwartzer et al., 2013; Smith et al., 2007), prenatal poly (I:C) injection on gestational day 12.5 reduced social preference for a novel same-sex individual. We speculate that for our MIA-treated SD animals, interactions with novel mice were experienced as a stressor rather than as an appetitive experience. Consistent with this interpretation, many of our SD-poly (I:C) animals had negative social preference scores suggesting that, in addition to disrupted social motivation, these animals had increased levels of social aversion as well. This notion is further supported by data of MIA mice engaging in higher rates of unsupported rearing behaviors during the social preference test.

Unsupported rearing is a type of risk assessment behavior displayed when confronted with threatening stimuli (Blanchard et al., 2011; Blanchard and Blanchard, 1989). Stressful experiences also promote the expression of this behavior (Sturman et al., 2018). Together, the reduced social preference and the enhanced display of social aversion and rearing are indicative of heightened stress reactivity in SD-poly (I:C) mice. As we have found previously with the LPS-MIA model in rats (Connors et al., 2014; Núñez Estevez et al., 2020; Zhao et al., 2020) life-long EE housing prevented reductions in social interaction and seemed to attenuate activation of the stress response.

### The Effects of MIA on Hippocampal Activity and the Display of Social Behavior

MIA may also impair approach-avoidance conflict decision-making, implicating the hippocampus in disrupted social interactions. The decision to approach or to avoid a novel social stimulus is likely the net result of multiple (conflicting) motives including the motivation to approach an appetitive stimulus and the motivation to avoid a potential threat (Brodkin et al., 2004; Elliot and Covington, 2001). Aberrant approach/avoidance processes are linked to autism, depression, and anxiety disorders (Heuer et al., 2007; Pfaff and Barbas, 2019; Radke et al., 2014). In an anxiety-provoking condition, SD-poly (I:C) mice may be less efficient at solving approach-avoidance decision-making conflicts. This notion is supported by our finding of diminished *c-Fos* immunoreactivity in the dentate gyrus, a region implicated in the regulation of approach-avoidance behaviors under innately anxiogenic and stressful conditions (Kheirbek et al., 2013; Weeden et al., 2015). However, data are inconsistent regarding the direction of dentate gyrus’ effects on approach behaviors (Kheirbek et al., 2013; Weeden et al., 2015). Although the specific role of the dentate gyrus is beyond the scope of the current study, our results suggest that reduced dentate gyrus activity may contribute to the impaired social preference in SD-MIA mice. While EE housing did not prevent this reduced neural activity following the social exposure, its beneficial effects appear to be mediated through other mechanisms more specific to HPA axis regulation.

### The Effects of MIA and EE on Hippocampal Feedback Regulation of the HPA Axis

In general, MIA appears to disrupt the hippocampal feedback regulation of the HPA axis, particularly in males, leading to delayed stress recovery following social stressor exposure. This is supported by the ventral hippocampal mRNA expression data showing higher levels of Crh in both male and female SD-poly (I:C) mice, 90 minutes following stress, compared to the SD-saline controls. Also, CRH and glucocorticoid receptors (Crhr1 and Nr3c1), as well as plasma corticosterone, were elevated in SD-poly (I:C) males, underscoring the sex-specific nature of this HPA axis dysregulation, despite the shared behavioral phenotypes. The occurrence of males and females displaying similar behavioral phenotypes regulated through separate sex-specific mechanisms is not unique (see Sorge et al., 2015). Additionally, there are well known sex differences that underlie the function of the stress axis. For example, sex differences in Crhr1 receptor binding may account for differences in receptor number and distribution (Kokras et al., 2019). It is likely that the HPA axis is driving some of the behavioral phenotypes that follow MIA, at least in part, but the female biomarkers only sometimes align with males due to the sex differences that underlie these mechanisms.

In addition to sex differences, there are also housing differences in HPA axis regulation. While EE attenuated many of the hippocampal gene expression changes that accompanied MIA, curiously, Crh expression and plasma corticosterone was elevated in EE-saline mice. This could be indicative of the ‘double edged sword’ of EE, in that for some male animals, it may be aversive (McQuaid et al., 2012; McQuaid et al., 2013). Especially with male CD-1 mice, EE can induce fighting depending on the enrichment devices used. However, in our C57BL/6J strain we did not observe physical indicators of distress or fighting among animals. One consideration is that we did not measure Crhr2, and the relative expression of these receptors, alongside Crhr1 (Skelton et al., 2000; Muller et al., 2003; Bale & Vale, 2004; Greetfeld et al., 2009; Wang et al., 2012), may be responsible for the more regulated social interactions and stressor responses demonstrated by EE-saline mice. Indeed, in our previous work we have shown EE housing to decrease the relative Crhr1/Crhr2 expression in the brain (Kentner et al., 2018c). Moreover, receptor expression and engagement with their ligands are what underlie neurophysiological and behavioral phenotypes.

Overall, it is unclear whether EE-saline males were displaying indicators of a disrupted stress recovery. However, elevated hippocampal glucocorticoid receptor expression (Vivinetto et al., 2013) and baseline corticosterone (Konkle et al., 2010; Moncek et al., 2004) have been reported in EE housed animals previously; the latter hypothesized as a potential indicator for eustress (Selye, 1956; Konkle et al., 2010). It is possible that poly (I:C) impeded these effects through separate mechanisms. However, given the protection of the social behavior phenotypes, it could be that EE overcompensated against the effects of MIA. Since EE-saline mice did not demonstrate behavioral impairments we do not believe that the elevated HPA activity in these animals are indicative of pathogenic processes.

### Novel Sex-Specific Effects of MIA on the Oxytocin System in the Ventral Hippocampus

The antistress effects of oxytocin in the hippocampus (Matsuchita et al., 2019) are altered by MIA challenge. In this study we observed elevations of hippocampal Oxtr in MIA males. A similar effect was observed in adult male rats that underwent prenatal restraint stress; these animals displayed an anxious/depressive phenotype, alongside reductions in the duration of social interactions and depolarization-evoked glutamate release in the ventral hippocampus (Mairesse et al., 2015). Other work has also shown early-life stress to alter the development of hippocampal oxytocin receptors (Noonan et al., 1994). Moreover, chronic stress and corticosterone implants increase oxytocin receptor binding in the ventral hippocampus (Liberzon and Young, 1997). However, the specific roles of increased oxytocin receptor binding remain unclear.

To our knowledge, this is the first study revealing effects of MIA and EE on the ventral hippocampal oxytocin system in females. Notably, there are significant sex differences in oxytocin receptor binding throughout forebrain regions of the brain that correlate with social interest (Dumais et al., 2013). Indeed, *upregulation* of Oxtr mRNA in area CA1 of the female hippocampus is hypothesized to increase social interest (Quiñones-Jenab, 1997; Dumais et al., 2013). Here, we observed reduced hippocampal Oxtr and social interest in our SD-poly (I:C) female mice, an effect rescued by EE. Overall, our results suggest opposite effects on Oxtr expression in male and female despite both sexes demonstrating similar social phenotypes. This again suggests that females are vulnerable to MIA challenges, but their impairments are at least in part mediated through separate mechanisms from males (Núñez Estevez et al., 2020).

## Conclusions

### The Protective Potential of Supportive Interventions

The current study demonstrates that the disruptive effects of MIA on social behaviors are associated with HPA axis dysregulation in the ventral hippocampus. The protective effects of EE reported here are consistent with our previous studies showing that EE can rescue MIA-impaired social behaviors induced by LPS (Connors et al., 2014; Kentner et al., 2016; Kentner et al., 2018b; Núñez Estevez et al., 2020; Zhao et al., 2020). Life-long EE may serve as an intervention to rescue these deficits by its ability to dampen activity of the HPA axis. When interpreting these findings, it is important to note that the validation of our model was done by evaluating maternal body weight gain following poly (I:C) challenge. The lack of maternal cytokine data to accompany this validation protocol is a limitation of this work. Additionally, it is difficult to isolate the critical timing of EE exposure that produced the beneficial effects reported in this study, and whether they are due to enhanced parental care during the early neonatal period. This is because EE removal induces depression-like behaviors and decreases the peak glucocorticoid response to acute stress (Smith et al., 2017). It should be noted that EE did not buffer all the MIA-induced changes in brain and behavior, but its success does appear to be most potent against social impairments and neural indicators of stress dysregulation.

Given the translational applications of EE as an intervention for humans (Janssen et al., 2014; Morgan et al., 2014; Rosbergen et al., 2017; Woo et al., 2015), our findings shed light on the sex-specific neural mechanisms that underlie its therapeutic success. Importantly, this work provides evidence for EE to be a viable supportive intervention against MIA in clinical settings, to be used alone or in combination with other treatments as appropriate.

## Funding and Disclosure

This project was funded by NIMH under Award Number R15MH114035 (to ACK) and the MCPHS Center for Undergraduate Research Mini-Grants (to RM & HT). The authors wish to thank Ryland Roderick and Madeline Puracchio for technical support during the early phases of this study. The authors would also like to thank the MCPHS University School of Pharmacy and School of Arts & Sciences for their continual support. The content is solely the responsibility of the authors and does not necessarily represent the official views of any of the financial supporters.

## Author Contributions

X.Z., R.M., H.T., M.E., & A.C.K., ran the experiments; X.Z., & A.C.K. analyzed and interpreted the data, and wrote the manuscript; A.C.K., designed, supervised the study, and made the figures.

## Declaration of Competing Interest

The authors declare that they have no known competing financial interests or personal relationships that could have appeared to influence the work reported in this paper.

## Notes

### Competing Interest Statement

The authors have declared no competing interest.

### Summary of Updates

updated figures and revised manuscript (primarily discussion section)

