## Supplementary Materials - Supplementary Fig S1 and Supplementary Tables S2 & S3 for "Poly (I:C)-induced maternal immune activation modifies ventral hippocampal regulation of stress reactivity: prevention by environmental enrichment"

Supplementary Figure S1
Supplementary Table S2
Supplementary Table S3

**Supplementary Figure S1**. **The effects of maternal immune activation (MIA) and environmental enrichment on P70 offspring open field and P85 social investigation rearing and grooming behaviors**. Postnatal day (P) 70 open field data on the number of rears for **(A)** male and **(B)** female mice. P85 data on the number of rears for **(C)** male and **(D)** female mice during the social exposure. Grooming behavior (seconds) in **(E)** males and **(F)** females during the P70 open field test and in **(G)** males and **(H)** females during the P85 social exposure. Data are expressed as Mean ± SEM, n=8-14 litters represented per sex, MIA, and housing group. $^{a}p < 0.05$, main effect of housing; $^{c}p < 0.05$, main effect of sex.

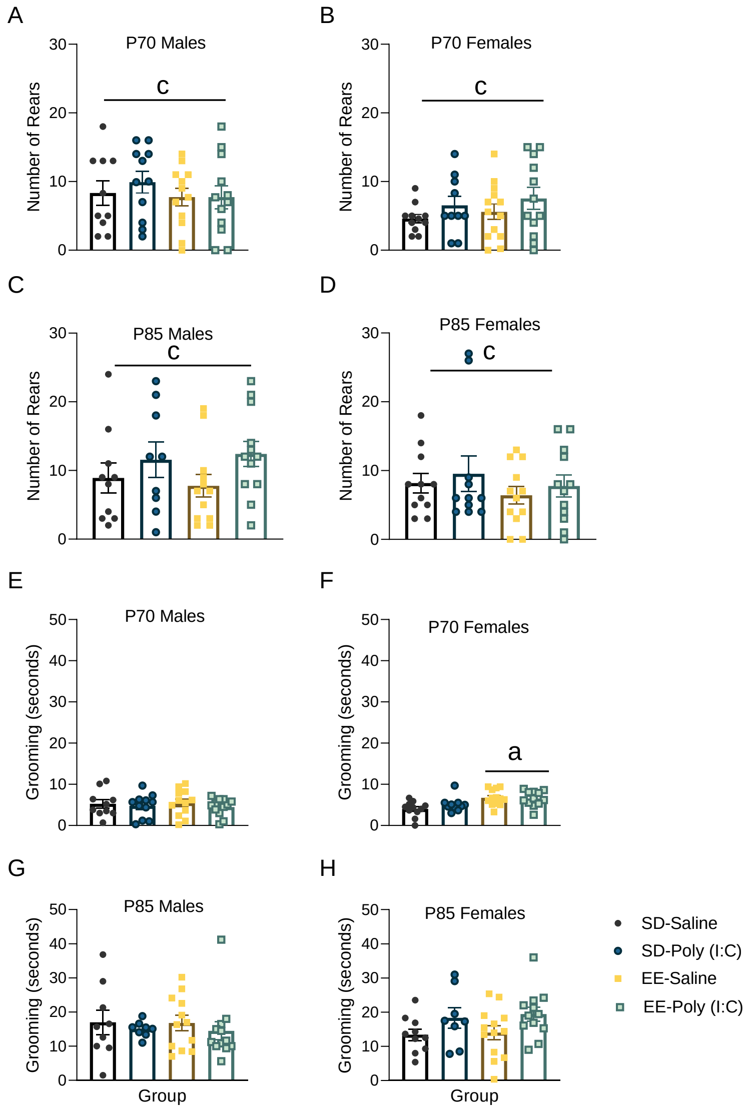


**Supplementary Table S2. Litter Designations and Treatments.**

| Number of Dams Included in Maternal Measures | Number of Offspring Included in P70 Behavioral Analyses | Number of Offspring Included in P85 Behavioral Analyses | Number of Offspring Included in qPCR Analyses | Number of Offspring Included in cFos Analyses |
|---|---|---|---|---|
| SD-Saline; (n = 12)[†] | 11 Males; 10 Females[#] | 9 Males[#] ; 11 Females | 8 Males; 8 Females | 7-8 Males; 9 Females |
| SD-Poly (I:C); (n = 14)[†] | 11 Males[*#]; 10 Females[*] | 12 Males[*]; 10 Females[*] | 8 Males; 8 Females | 7-8 Males; 7-8 Females |
| EE-Saline; (n = 12) | 12 Males; 13 Females[*] | 10 Males[#]; 13 Females[*] | 8 Males; 8 Females | 7-8 Males; 6-7 Females |
| EE-Poly (I:C); (n = 12) | 12 Males; 12 Females | 12 Males; 12 Females | 8 Males; 8 Females | 7-8 Males; 7-8 Females |
| N = 50 dams; N = 93 offspring | N = 91 | N = 89 | N = 64 | N = 58-64 |

Note: 65 dams were bred. 8 dams lost their litters during pregnancy or had small litters (n < 4 pups) that were fostered to same age and group matched dams. These issues were primarily in the SD-Poly (I:C) group. 7 dams did not become pregnant.
#Animal not evaluated on behavioral measure due to experimenter error or video recording failure.
† Litter lost after birth.
*Extra pups in litter represent fostered animals.

**Supplementary Table S3. mRNA gene expression in offspring exposed to maternal immune activation and housed in SD or EE**

| Brain region | Sex | Neural marker | SD | | EE | |
|---|---|---|---|---|---|---|
| | | | Saline | Poly (I:C) | Saline | Poly (I:C) |
| Prefrontal Cortex | *Males* | *Crh* [**] | 1.00 ± 0.12 | 1.64 ± 0.30 | 0.85 ± 0.08 | 1.30 ± 0.28 |
| | | *Crhr1* [**] | 1.00 ± 0.08 | 1.30 ± 0.11 | 1.17 ± 0.11 | 1.39 ± 0.05 |
| | | *Nr3c1* [##] | 1.00 ± 0.09 | 1.36 ± 0.13 | 0.76 ± 0.08 | 0.96 ± 0.16 |
| | | *Oxt* [†] | 1.00 ± 0.05 | 1.07 ± 0.34 | 0.56 ± 0.18[a] | 1.08 ± 0.26 |
| | | *Avpr1a* | 1.00 ± 0.63 | 2.85 ± 1.10 | 1.56 ± 0.61 | 1.66 ± 0.57 |
| | | *Oxytr* | 1.00 ± 0.15 | 1.31 ± 0.18 | 0.97 ± 0.20 | 1.04 ± 0.13 |
| | *Females* | *Crh* | 1.00 ± 0.20 | 0.86 ± 0.07 | 0.77 ± 0.07 | 0.73 ± 0.11 |
| | | *Crhr1* [** ##] | 1.00 ± 0.10 | 0.89 ± 0.04 | 1.51 ± 0.04 | 1.13 ± 0.07 |
| | | *Oxt* | 1.00 ± 0.34 | 0.63 ± 0.19 | 0.76 ± 0.32 | 0.44 ± 0.13 |
| | | *Avpr1a* | 1.00 ± 0.27 | 1.37 ± 0.36 | 0.97 ± 0.374 | 1.35 ± 0.40 |
| | | *Oxytr* | 1.00 ± 0.18 | 0.68 ± 0.06 | 0.84 ± 0.14 | 0.62 ± 0.06 |
| | | *Nr3c1* [##] | 1.00 ± 0.09 | 1.15 ± 0.07 | 0.84 ± 0.07 | 0.80 ± 0.12 |
| Ventral Hippocampus | *Males* | *Crh* [††] | 1.00 ± 0.08 | 2.71 ± 0.397[ab] | 2.00 ± 0.27[ab] | 1.18 ± 0.14 |
| | | *Crhr1* [††] | 1.00 ± 0.10 | 1.69 ± 0.116[ab] | 0.96 ± 0.17 | 0.85 ± 0.17 |
| | | *Nr3c1* [†††] | 1.00 ± 0.05 | 2.88 ± 0.16[ab] | 1.69 ± 0.10[a] | 1.32 ± 0.26 |
| | | *Oxt* | 1.00 ± 0.45 | 0.88 ± 0.12 | 1.01 ± 0.35 | 0.89 ± 0.22 |
| | | *Avpr1a* [†] | 1.00 ± 0.20 | 2.05 ± 0.38[a] | 2.25 ± 0.58 | 1.70 ± 0.26 |
| | | *Oxytr* [††] | 1.00 ± 0.12 | 2.13 ± 0.29[ab] | 1.37 ± 0.27 | 1.34 ± 0.19 |
| | | *Camk2a* [†] | 1.00 ± 0.09 | 1.68 ± 0.16[ab] | 1.07 ± 0.06 | 1.24 ± 0.11 |
| | | *Prkca* [††] | 1.00 ± 0.05 | 2.91 ± 0.19[ab] | 1.79 ± 0.15[a] | 1.44 ± 0.28 |
| | *Females* | *Crh* [†] | 1.00 ± 0.21 | 1.36 ± 0.07[b] | 0.84 ± 0.11 | 0.61 ± 0.05 |
| | | *Crhr1* | 1.00 ± 0.10 | 1.68 ± 0.32 | 1.61 ± 0.36 | 1.75 ± 0.19 |
| | | *Nr3c1* | 1.00 ± 0.20 | 1.13 ± 0.10 | 0.94 ± 0.08 | 0.63 ± 0.03 |
| | | *Oxt* | 1.00 ± 0.18 | 0.84 ± 0.18 | 0.61 ± 0.15 | 0.40 ± 0.07 |
| | | *Avpr1a* | 1.00 ± 0.26 | 1.75 ± 0.46 | 1.18 ± 0.11 | 1.04 ± 0.20 |
| | | *Oxytr* [††] | 1.00 ± 0.15 | 0.61 ± 0.06[ab] | 0.76 ± 0.10 | 0.97 ± 0.10 |
| | | *Camk2a* | 1.00 ± 0.13 | 1.11 ± 0.09 | 1.22 ± 0.07 | 1.02 ± 0.12 |
| | | *Prkca* [†] | 1.00 ± 0.19 | 1.39 ± 0.13[b] | 0.99 ± 0.07[ab] | 0.70 ± 0.08 |
| Hypothalamus | *Males* | *Crh* | 1.00 ± 0.14 | 1.01 ± 0.43 | 1.24 ± 0.63 | 1.32 ± 0.45 |
| | | *Crhr1* | 1.00 ± 0.24 | 1.49 ± 0.54 | 0.49 ± 0.07 | 1.44 ± 0.30 |
| | | *Nr3c1* | 1.00 ± 0.27 | 2.18 ± 0.80 | 2.08 ± 0.83 | 1.36 ± 0.28 |
| | | *Oxt* | 1.00 ± 0.18 | 1.98 ± 0.95 | 2.86 ± 1.86 | 1.93 ± 0.47 |
| | | *Avpr1a* | 1.00 ± 0.13 | 0.51 ± 0.30 | 0.97 ± 0.42 | 1.02 ± 0.35 |
| | | *Oxytr* | 1.00 ± 0.22 | 1.41 ± 0.28 | 1.12 ± 0.25 | 1.20 ± 0.25 |
| | *Females* | *Crh* | 1.00 ± 0.15 | 4.52 ± 2.02 | 2.91 ± 1.32 | 1.25 ± 0.24 |
| | | *Crhr1* | 1.00 ± 0.63 | 0.25 ± 0.07 | 0.15 ± 0.09 | 0.25 ± 0.05 |
| | | *Nr3c1* | 1.00 ± 0.26 | 1.60 ± 0.56 | 1.64 ± 0.85 | 0.77 ± 0.19 |
| | | *Oxt* | 1.00 ± 0.17 | 1.03 ± 0.23 | 1.63 ± 0.55 | 1.54 ± 0.52 |
| | | *Avpr1a* | 1.00 ± 0.16 | 2.01 ± 0.53 | 1.34 ± 0.47 | 0.92 ± 0.16 |
| | | *Oxytr* | 1.00 ± 0.17 | 0.58 ± 0.11 | 0.86 ± 0.22 | 0.93 ± 0.15 |

Data are mean ± SEM.

† P<0.05, significant interaction (MIA x Housing)

∗ P<0.05, ** P<0.01, significant main effect of MIA (saline vs. poly (I:C))

### P<0.05, ## P<0.01, significant main effect of housing (SD vs. EE)

Lower case letters indicate significant differences between conditions ([a]different from SD-Saline, [b]different from EE-Poly (I:C))
